# Efficient Spline Regression for Neural Spiking Data

**DOI:** 10.1101/2020.09.01.276105

**Authors:** Mehrad Sarmashghi, Shantanu P Jadhav, Uri Eden

## Abstract

Point process generalized linear models (GLMs) provide a powerful tool for characterizing the coding properties of neural populations. Spline basis functions are often used in point process GLMs, when the relationship between the spiking and driving signals are nonlinear, but common choices for the structure of these spline bases often lead to loss of statistical power and numerical instability when the signals that influence spiking are bounded above or below. In particular, history dependent spike train models often suffer these issues at times immediately following a previous spike. This can make inferences related to refractoriness and bursting activity more challenging. Here, we propose a modified set of spline basis functions that assumes a flat derivative at the endpoints and show that this limits the uncertainty and numerical issues associated with cardinal splines. We illustrate the application of this modified basis to the problem of simultaneously estimating the place field and history dependent properties of a set of neurons from the CA1 region of rat hippocampus, and compare it with the other commonly used basis functions. We have made code available in MATLAB to implement spike train regression using these modified basis functions.

## Introduction

Statistical neural models are often used to relate the likelihood of observing particular spike patterns in individual neurons to a variety of factors, including biological and behavioral signals, the neuron’s own past spiking history, and the influences of other neurons in a local population [1]. For instance, statistical models for neurons in the CA1 region of rat hippocampus have been used to model these neurons’ place field properties, theta rhythmicity and precession [2, 3], and the neuron’s past spiking history [4]. In order to understand neural receptive fields, point process generalized linear models (GLMs) are often used [1, 5–8]. In general, GLMs expand upon classic linear models to allow for a broad range of probability models including point process distributions, which are most appropriate for analysing neural spiking. GLMs provide robust and computationally efficient methods for estimating model parameters. They also provide powerful analysis tools for computing model uncertainty, assessing goodness-of-fit, and model refinement [1, 9–12]. Many researchers have applied point process GLMs to understand the coding properties of individual neurons in neural populations [1, 8, 13–16], other researchers have used them to infer biophysical features of neural systems [17–23], yet other analyses use GLMs to infer the dynamics of the unobserved underlying signals driving neural systems [24–27].

Often, point process GLMs are constructed using basis functions to capture complex functional relationships between neural spiking and the signals that influence it. These signals can include biological and behavioral signals, the activity of other neurons, or a neuron’s own past spiking history. In particular, splines are a common choice of basis functions that use locally defined, 3rd order polynomials to capture nonlinear relationships between spiking and covariates. Splines are flexible, have parameters that are interpretable, and have been successful in capturing nonlinear functional relationships in a variety of neural systems [2, 8, 28]. The most common class of splines used for point process neural modeling are cardinal splines. Each cardinal spline is defined by a sparse set of knots, or control points, such that the value of the function at any point along the spline and the value of its derivative are determined by a linear combination of the four nearest control points [29, 30].

Cardinal splines have been successfully deployed to capture nonlinear influences on neural spiking activity, however there are a number of issues that can arise when cardinal splines are used to model the influence of signals that are bounded above or below. This occurs, for example, when modeling hippocampal place fields in a bounded environment. Another example is modeling history dependent spiking, where a neuron’s recent spiking activity can influence current spiking; only spikes that occur before the current time can have an influence on current spiking, placing a bound on the temporal influences of spiking. One issue with standard cardinal spline basis functions is a loss of statistical power in estimating the influence of a covariate near its bounds relative to estimating the influence away from the bound. This is a natural consequence of having samples observed only on one side of each bound, but is exacerbated by the need to estimate the derivative of the spline at the bounds with limited data. In cases where firing rates are suppressed when a covariate approaches its bound, such as with refractoriness in history dependent models, cardinal splines can often lead to numerical instability in estimating the model parameters. Even with high firing rates near the bounds, this loss of power leads to large confidence intervals. Finally, limited data near the bounds can lead to estimates of the derivative of the spline function with large magnitude, that suggest rapid changes in firing responses that may not capture known receptive field structure. Together, these issues can make it difficult to assess the significance of the influence of bounded neural signals near their bounds.

The primary purpose of this paper is to illustrate the challenges associated with estimating uncertainty in point process neural receptive field models near the boundary of a bounded predictor and to present one option for meeting these challenges. In this work, we develop a simple modification to the cardinal spline basis functions to allow us to account more appropriately for sampling at the boundary. The issues discussed above are related to the fact that cardinal spline regression models use the data to estimate the intensity of spiking activity and its derivative simultaneously. These issues may be assuaged by making an assumption about the derivative near the boundaries to preserve statistical power. We construct a modified cardinal spline basis, assuming that its derivative is zero at the boundary. This reduces the number of parameters that need to be estimated, alleviates some of the unnecessary reduction in statistical power near the boundary, and leads to more computationally robust model fits. We illustrate the application of this model to the problem of simultaneously estimating the place field and history-dependent properties of 45 neurons recorded from the CA1 region of rat hippocampus [15].

The remainder of this paper is structured as follows. First, we review the fundamentals of point process generalized linear modeling (GLM) and its application to neural coding problems. Then, we explain the derivation of cardinal splines and their ability to capture nonlinear functional relationships. Next, we derive the modified cardinal spline basis functions. Finally, we apply these modified basis functions to real data recorded from the rat hippocampus and demonstrate how they can limit the uncertainty and numerical issues at the boundaries. We also compare the modified splines with the other commonly used basis functions in order to understand their properties.

## Materials and methods

We begin by reviewing the point process GLM modeling framework for neural spike train data along with the definition of cardinal splines and their use as basis functions in this framework. Next, we propose a modification to these functions that improve statistical power and limit numerical instability near the boundaries of the factors influencing spiking. Finally, we discuss maximum likelihood parameter estimation and methods for evaluating the goodness-of-fit of these models to the data.

### Point Process-GLM Framework

A point process provides a probability model for a set of binary events occurring in continuous time. Point process models are often used to characterize the factors that influence spiking activity in individual neurons or neural populations [1]. Any point process can be characterized by its conditional intensity function [31],

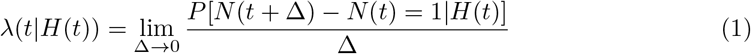

where *N* (*t*) is number of spikes in time interval (0, *t*] and *H*(*t*) is the past spiking history of the neuron or neural population being modeled up to time *t*. For small Δ, *λ*(*t*| *H*(*t*))Δ is approximately the probability of observing a spike in the time interval (*t, t* + Δ] given the spiking history.

A neural point process model expresses *λ*(*t*|*H*(*t*)) as a function of a set of covariates thought to influence spiking activity. Point process GLMs are one class of models that are easy to fit, interpret, and assess. These models relate the firing intensity to a linear combination of functions of the covariates influencing spiking through a nonlinear link function. One common GLM structure defines the log of the conditional intensity of each neuron to be linear combinations of functions of extrinsic covariates related to spiking, functions of the neuron’s own spike history, and functions of the past spiking activity of other, simultaneously recorded neurons [1]

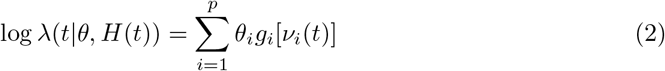

Where *g*_*i*_(·) is a basis function acting on a covariate *ν*_*i*_(*t*), and *p* is the dimension of the model parameter *θ*.

GLMs allow researchers to capture nonlinear, complex relationships between stochastic signals in a manner that is flexible, computationally efficient and robust. They also provide powerful analysis tools for computing model uncertainty, assessing goodness-of-fit, and refining models [1, 9–11]. In addition, for the GLM expressed in Eq (2) it is easy to prove that the log-likelihood function of the data is guaranteed to be concave as a function of the model parameters, which means that maximum likelihood estimation is guaranteed to converge to the global maximum [1]. These models are a generalized form of linear models so they are intuitive and readily interpretable. However, GLMs assume that the data comes from a specific set of distributions known as the natural exponential family. The degree of fit between the data and the assumed distributional model must be verified. Fortunately, GLMs are equipped with a number of goodness-of-fit tools to assess those assumptions [32, 33]. In addition, while GLMs are flexible and able to capture arbitrary non-linear and high-dimensional relationships, this flexibility may require a large number of parameters to estimate, which can lead to a trade-off between model size and accuracy, and make estimating high-dimensional relationships impractical. Moreover, a number of researchers have compared the success of GLMs against other model structures including modern machine learning algorithms, and found situations in which those models outperform GLMs [23, 34–38].

In the data analysis example below, we model the firing intensity of a population of neurons in the CA1 region of the hippocampus of rats performing a spatial navigation task by letting *ν*_*i*_(*t*) include both the rat’s location and the past spiking history of the modeled neuron. We will compare different sets of basis functions for these covariates.

### Spline Functions

Splines are piecewise, low-degree polynomials that interpolate continuously and smoothly between a set of control points. These functions were proposed to resolve the severe oscillation of high-degree polynomials in large intervals particularly (Runge’s phenomenon) [39]. There are multiple classes of splines, which reflect different choices in the degree of polynomials, the functions used to represent those polynomials, and the assumptions about these functions’ smoothness. A common choice for spike train models is the cardinal spline, which uses cubic Hermite polynomials to interpolate between a set of control points and has derivatives at each control point determined by neighboring control points (Fig 1B) [40]. To be clear, each local segment of the spline is a third order polynomial, but the spline function is not a third order polynomial; it is sufficiently flexible to approximate arbitrary smooth functions no matter what order. Typically, cubic cardinal splines, defined by third-order polynomials, are used for neural point process regression. If we let *x*_*i*_ represent the location of the *i*^*th*^ control point, the equation for the spline at a point *x* between control points *x*_*i*_ and *x*_*i*+1_ is

**Fig 1.**
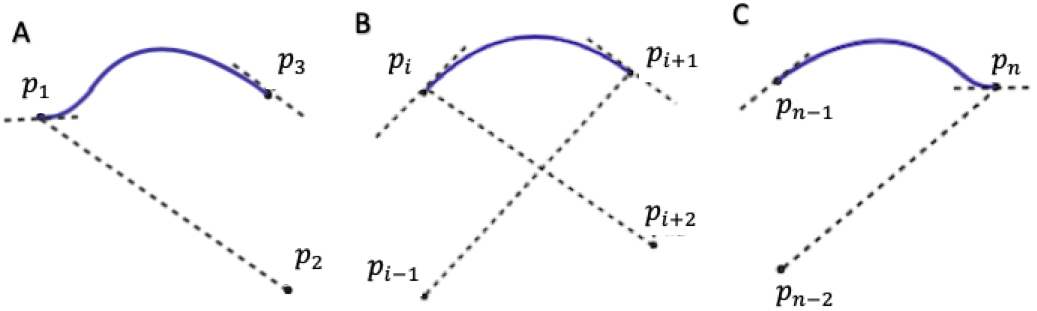
Modified cardinal spline function. Modified spline function at A: Initial segment, B: Middle segments, and C: Final segments.

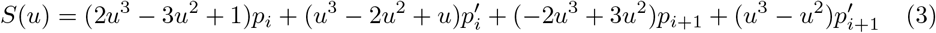

Here,

- 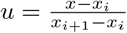
- *p*_*i*_ = *f* (*x*_*i*_) is the value of the function at the *i*^*th*^ control point and 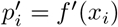

It is clear from Eq (3) that *S*(*u*) passes through the control points. The derivatives at the control points are determined by a tension parameter, *s*, and the values at neighboring control points:

- 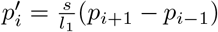 where 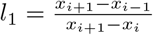
- 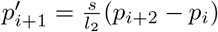 where 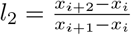

The general expression for the cardinal spline is therefore

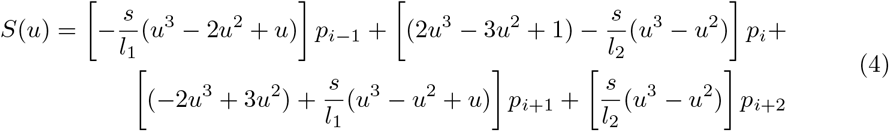

Eq (4) indicates that the value of a cardinal spline at any point is given by a linear combination of four nearest control points. Standard cardinal splines are not defined at the segments between the first and second control point or between the second-to-last and last control point. They are only defined at the locations for which two control points to the left and two control points to the right exist. At the boundary, modelers typically place two control points beyond the most extreme value of the covariate. The first and last control point therefore do not reflect the values of the functional relationship at their knot locations. They are only used to estimate the derivative at the boundary.

As discussed above, while these functions have been successfully used to model neural spiking activity, the specification of the derivative using surrounding control points can lead to loss of statistical power at the boundaries of *x*.

### Modified Cardinal Spline

In this section, we derive a modified spline function that assumes that the derivative is equal to zero at the end control points but uses the neighboring control points to determine the derivatives at interior control points (See Fig 1).

For any value of *x* such that there are at least two control points less than *x* and 2 control points greater than *x* (Fig 1 B), the spline equation is still given by Eq (4). We can also express Eq (4) in matrix form as follows:

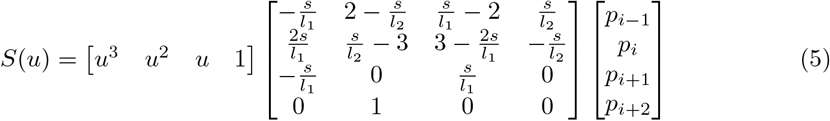

Near the lower boundary, for any value of *x* that has only one control point less than *x* (but still two control points greater than *x*), we construct a spline segment satisfying these conditions (Fig 1A):

- *S*(*u*)|_*u*=0_ = *p*_1_ and *S*(*u*)|_*u*=1_ = *p*_2_
- *S*′(*u*)|*u*=0 = 0 and 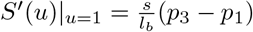 where 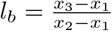

Plugging these conditions into Eq (3) we obtain the following solution for the initial spline segment:

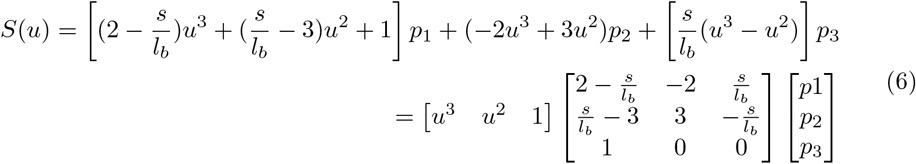

Near the upper boundary, for any value of *x* such that there is only one control point greater than *x* (but there are still two control points less than *x*) we construct a spline segment satisfying these conditions (Fig 1C):

- *S*(*u*)|_*u*=0_ = *p*_*n*−1_ and *S*(*u*)|_*u*=1_ = *p*_*n*_
- 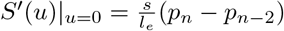 where 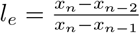 and *S*′(*u*)|_*u*=1_ = 0

Here, *n* is the total number of control points. Plugging these conditions into Eq (3) we obtain the following solution for the final spline segment:

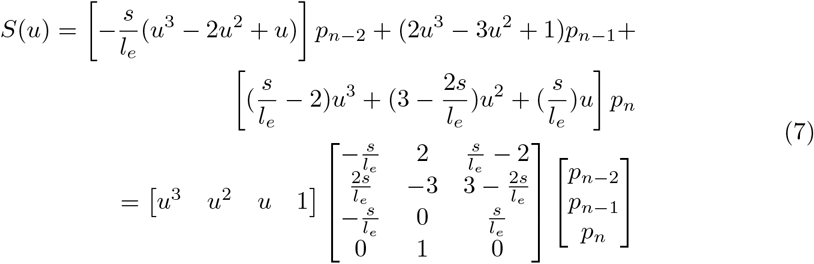

The above expressions fully define the modified cardinal spline in the case where at least three control points are used. In the unusual case where a model is built using a spline with only two control points, we can solve for the single segment between these control points based on the following conditions

- *S*(*u*)|_*u*=0_ = *p*_1_ and *S*(*u*)|_*u*=1_ = *p*_2_
- *S*′(*u*)|_*u*=0_ = *S*′(*u*)|_*u*=1_ = 0

Plugging these conditions into Eq (3) we obtain the following solution for a model with a single spline segment:

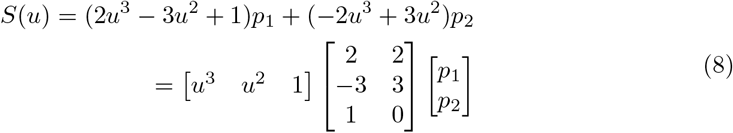

### Maximum Likelihood Parameter Estimation

Point process GLMs of the form of Eq (2) have a number of properties that make it easy to estimate model parameters, construct confidence intervals, and assess goodness-of-fit. The form of this model guarantees that the likelihood surface is convex, ensuring that there exists a unique maximum. This is true even when the functional relationship expressed in right hand side of Eq (2) includes flexible, non-convex functions of the covariates. For GLMs, this maximum is typically found using an iteratively re-weighted least squares (IRLS) procedure, due to its computational simplicity, efficiency and robustness [1]. In addition to finding the maximum likelihood estimates of all of the model parameters, the IRLS procedure provides the observed Fisher information matrix, 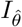, which allows for the construction of confidence intervals and standard tests of significance of the influence of individual or multiple covariates [1, 10, 11]. The maximum likelihood estimates for the parameters multiplying the modified spline basis functions are directly interpretable; since the spline interpolates between the control points, when the signal, *x* takes a value equal to the control point location associated with basis element *g*_*i*_, the conditional intensity is equal to 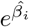. Confidence intervals for each spline parameter can be computed directly from the diagonal entries of inverse of the Fisher information, 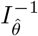. The intensity and confidence associated with other values of *x* can also be computed, but require incorporating the estimated covariance between multiple parameter estimates using the full Fisher information. Statistical software packages such as R and MATLAB include routines that can be used to fit point process GLMs and compute confidence bounds; these routines also output multiple goodness-of-fit measures, including the model deviance and model residuals.

### Data Analysis Example

To demonstrate the properties of our modified cardinal spline basis functions, we constructed history dependent place field models of a population of neurons in the CA1 region of rat hippocampus while the animal performed a spatial navigation task. One male Long–Evans rat weighing 450 –550 g was implanted with a movable array of recording tetrodes in the CA1 region of its hippocampus. The animal had been trained to perform a spatial alternation task on a W-shaped track, alternating between the left and right arms before returning to the center arm. It received a reward at the ends of the left and right arms after each correct alternation. The data used in this study is secondary data that has been reported previously on [15, 41], and is available publicly on CRCNS data sharing page.

We constructed point process GLMs for each of 45 neurons in this population based on Eq (2) as follows:

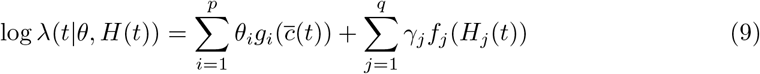

Here, the first sum represents the component related to the rat’s position; the second one represents the influence of the neuron’s own firing history. *θ*_*i*_, and *γ*_*j*_ are the model parameters. *g*_*i*_(·) represents a set of basis functions acting on the rat’s position 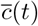 for a set of *i* = 1, 2, …, *p* parameters and *f*_*j*_(·) represents basis functions acting on the past spiking history of the neuron being modeled. We fixed the knot locations for each of the temporal and spatial components. For the unmodified cardinal spline we placed two control points beyond the most extreme value of the covariate. The first and last control point are only used to estimate the derivative at these locations.

For the specific model form expressed in Eq (9), we are assuming that the activity of each neuron is predicted only by position and the neuron’s own history and not by the activity of other neurons. We can further refine this model by including the activity of other neurons but we have deliberately chosen to fit a simple model in order to highlight issues related to uncertainty in point process GLM estimates near the boundary of bounded predictors.

We compared 4 choices of basis functions to capture the place fields and history dependence of these neurons: raised cosines, indicator functions, standard cardinal splines, and our modified splines. All of the models had a common form given by Eq (9), where *g*(·) and *f* (·) are the different choices of basis function for each model.

Raised Cosines are a set of restricted cosine functions with a logarithmic scaling of the *x* axis, given by the following expression

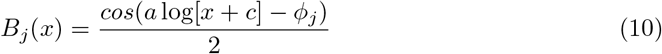

such that *a* log(*x* + *c*) ∈ [*phi*_*j*_ – π, *phi*_*j*_ + *π*] and 0 elsewhere. *a* and *c* are parameters that are set by hand and *ϕ*_*j*_ is a parameter that determines where the peaks occur with *π*/2 radian spacing between them. We construct raised cosine models by replacing the basis functions *g*(·) and *f* (·) in Eq (9) with *B*(·) from Eq (10). These bases allow for fine representation of time and space [42, 43].

Indicator Functions are a set of step functions that are nonzero over non-overlapping subsets of the domain of *x*. Here, the range of *x* is partitioned into a set of intervals, *A*_*j*_, and the indicator basis function are defined as:

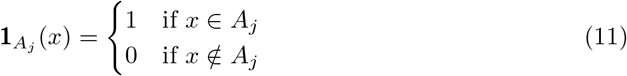

For a sufficiently small partition, these functions can capture rapid changes in the influence of *x* on the spiking intensity. We construct indicator models by using indicator functions as the bases *g*(·) and *f* (·) in Eq (9) to represent both the temporal and spatial components.

We compared the neural models in terms of the width of the confidence intervals for the history dependent component of the model. In order to quantify the effect of the choice of basis function on the size of the confidence bounds, we computed a confidence interval width ratio:

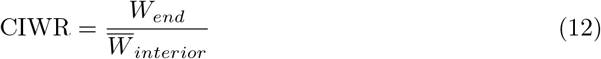

where *W*_*end*_ is the confidence interval width at the specified endpoint of the variable of interest, and 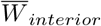 is the average of the confidence interval width in an interior region from 0.05 to 0.95 times the range of the variable.

The purpose of this comparison is not to argue that spline basis functions are generally superior to other basis functions for modeling neural activity. The choice of which basis functions to use depends primarily on the features of the problem to be addressed and the structure of the data. In this paper, the comparison to other commonly used basis functions is made primarily to provide intuition about the behavior of the modified spline at the boundary of bounded covariates, and how that compares with other common choices of basis functions.

### MATLAB Code

We provide software tools for constructing and fitting our modified spline basis functions using MATLAB on the companion GitHub repository. The repository contains functions for constructing modified spline basis functions (as well as the indicator, raised cosine, and unmodified spline bases) and code for fitting model parameters and computing confidence bound about individual parameters and the intensity function as a whole. For illustration purposes, we provide code that has been applied to simulated neural spiking data based on the features of our real data. It should be simple to adapt this code to any neural spiking data set.

## Results

It is well known that the activity of neural ensembles in the CA1 region of rat hippocampus correlates with the position of the animal when actively exploring an environment [2, 3] and with the past spiking history of each neuron [4]. Fig 2 shows an example of spatial and temporal coding of a single hippocampal neuron. Fig 2A shows the place field for the neuron with the blue trajectory showing the movement of the rat and the red dots showing the locations where the neuron fired. This neuron is more active when the rat is on the right side of the track. Fig 2B illustrates the place field structure of the neuron in a linearized version of this track, using an occupancy-normalized histogram. Each color represents the firing rate of the neuron in each arm of the maze. As we expect from Fig 2A, we observe the highest spiking activity in the right arm. Fig 2C shows the spike autocorrelation function, where the x-axis represents a time lag in ms. This neuron’s spiking is highly correlated with its past activity. The negative autocorrelation at a lag of 2 ms reflects the refractory period of the neuron. We have significant positive correlation at 3-70 ms lags which reflects the tendency of the neuron to fire with a theta rhythm. Fig 2D shows the interspike interval (ISI) histogram of the neuron. We observe a peak in ISIs between 5-25 ms interval, suggesting that the neuron tends to fire in bursts with short ISIs in its place field.

**Fig 2.**
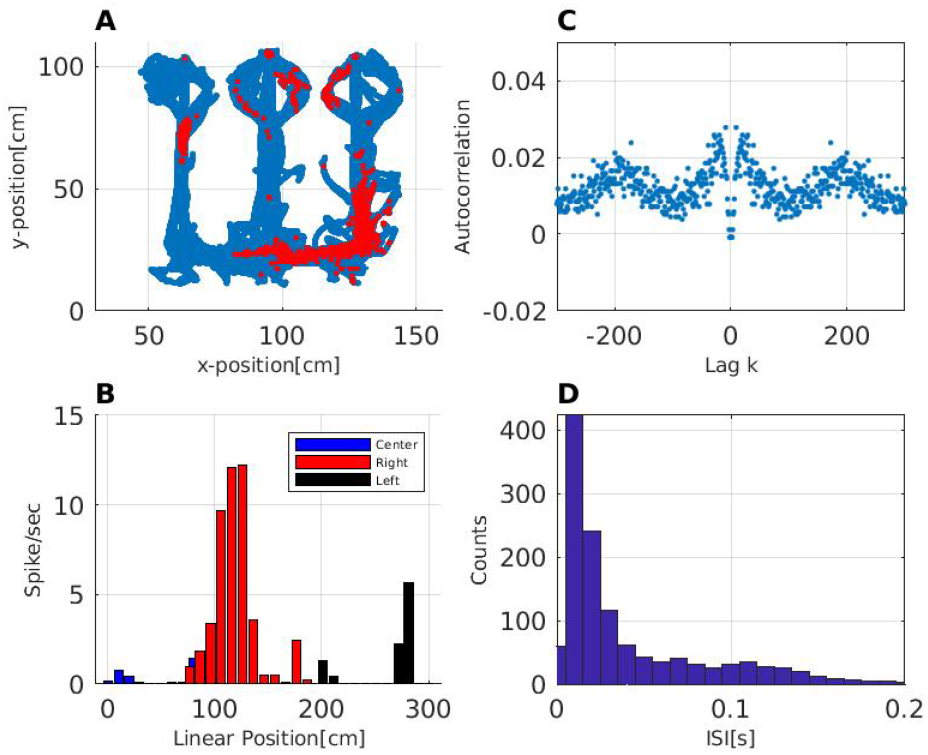
Activity of a single neuron in the hippocampus during behavior. A: Place field for one neuron. The rat’s movement trajectory is shown in blue and the locations where the neuron spikes is shown in red. B: Occupancy-normalized histogram of firing activity of the neuron in a linearized version of the environment. Each color represents the firing rate of the neuron in a different arm of the W-maze. C: Autocorrelation of the spike train of the neuron. The x-axis is time lag using 1 ms bins. D: Interspike interval (ISI) histogram.

We compared the properties of the modified spline to other basis functions often used to model spike trains: raised cosines, cardinal splines, and indicator functions. Fig 3 illustrates four example sets of bases functions used to model the dependence of neural spiking on its past history up to two hundred ms in the past. The selection of 200ms as the endpoint of the history dependence was based on both empirical evidence of the amount of prior history that contributes significantly to current spiking and to known physiological properties of these neurons such as bursting and theta rhythmicity. Here, our goal is to develop a causal models, hence the smallest possible time which one neuron can influence the next neuron is at 0 ms. If we did try to extend the control points to negative lag values, there would be no observed values of the predictors at negative value, thus at 0m there is an actual bound. However, there is no bound on 200, and it is possible to extend lags beyond it, because there exists spike data at these further lags. 50 indicator functions over each 4ms bin are shown in Fig 3A. Fig 3B shows a set of raised cosine basis functions, using a log-scaling of the time axis to provide fine representation at early lags and broader coverage at later lags. We have selected parameter values for indicator functions and raised cosines that are consistent with those typically used in the literature [27, 42, 43]. Fig 3C and 3D show classical and modified cardinal splines that also provide finer resolution at early lags. The modified spline uses two fewer basis functions and differs in the functions with support near the endpoints, but uses identical functions for the remaining basis elements.

**Fig 3.**
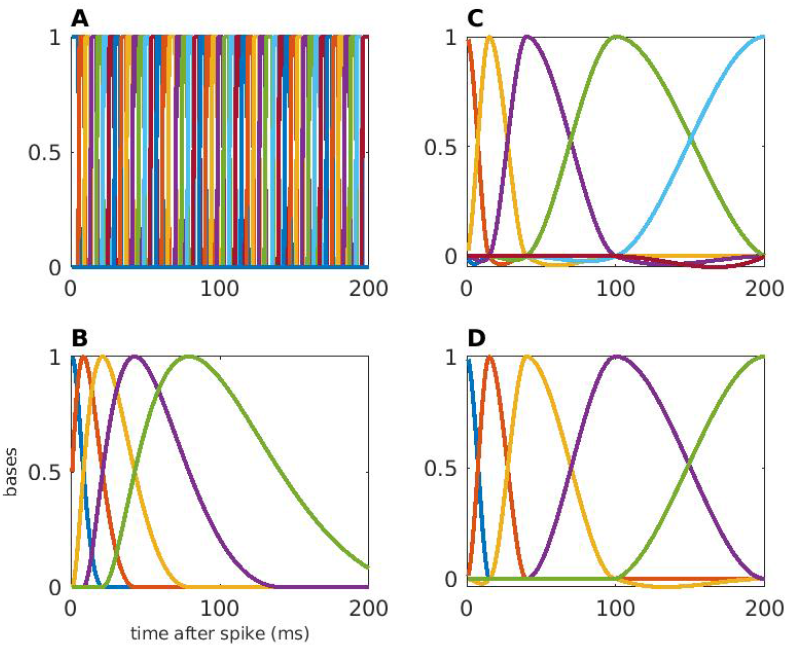
Sets of basis functions for comparison. We used four basis functions to represent spiking history A: 100-dimensional indicator function. B: 5-dimensional raised cosine function. C: 7-dimensional cardinal spline. D: 5-dimensional modified cardinal spline.

We used these bases in a point process GLM to capture the spatial and the temporal structure of the firing rate of each recorded neuron. Fig 4 shows an example of the fit for the spatial and history dependent components of the resulting model fits for a single neuron. Fig 4A shows the history dependent component going back two hundred lags in the past. All of the model fits capture the basic properties of refractoriness in the first few ms after a previous spike, and increased excitability around 20 after a previous spike. As expected, the model based on the indicator basis produces estimates that are less smooth than those based on the other 3. Additionally, the models differ in their estimated uncertainty, as evidenced by the size of their confidence bounds, especially near lags 0 and 200. For the model based on indicator basis functions, the variability of the estimates and large confidence bounds make it difficult to draw inferences about spiking dynamics from this model fit. For the model based on raised cosines, the size of the confidence intervals becomes smaller with increasing lag, to the point where they are very evidently too small at lags near 200, expressing unwarranted confidence based on the data. The logarithmic temporal scaling of the raised cosine functions means that estimates of history dependence at later lags integrate spiking over longer periods; a spike occurring 100 ms after another will influence the model estimate not only at a lag of 100 ms but also at a lag of 200 ms, leading to smoother estimates with smaller estimated uncertainty as lag increases. The computed CIWR value for the raised cosine model at lag 0 is 8.11 whereas it is 0.06 at lag 200, which corroborates the observation that the variability at later lags is underestimated. For the unmodified cardinal spline model, the confidence intervals have consistent width for lags in the interior, away from 0 and 200. Near lags 0 and 200 the magnitude of the confidence intervals increases substantially. Some of this is attributable to the fact that we have fewer spikes with which to estimate the parameters near the boundaries. We might expect the confidence bounds to increase by a factor of around 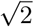 to account for the fact that about 1/2 the spikes are available for fitting these parameters. However, the estimated CIWR for the unmodified cardinal spline is 184.38 at lag 0 and 22.1 at lag 200, reflecting an extreme loss of certainty near the endpoints. The bottom right panel of Fig 4 shows how the modified cardinal spline basis can resolve this uncertainty and instability at the boundaries. The confidence intervals flare out slightly at the boundaries, reflecting the decrease in data for parameter estimation, but do not expand nearly as much as the unmodified spline model. The CIWR for the modified spline model is 2.93 and 1.54 at the beginning and end points respectively.

**Fig 4.**
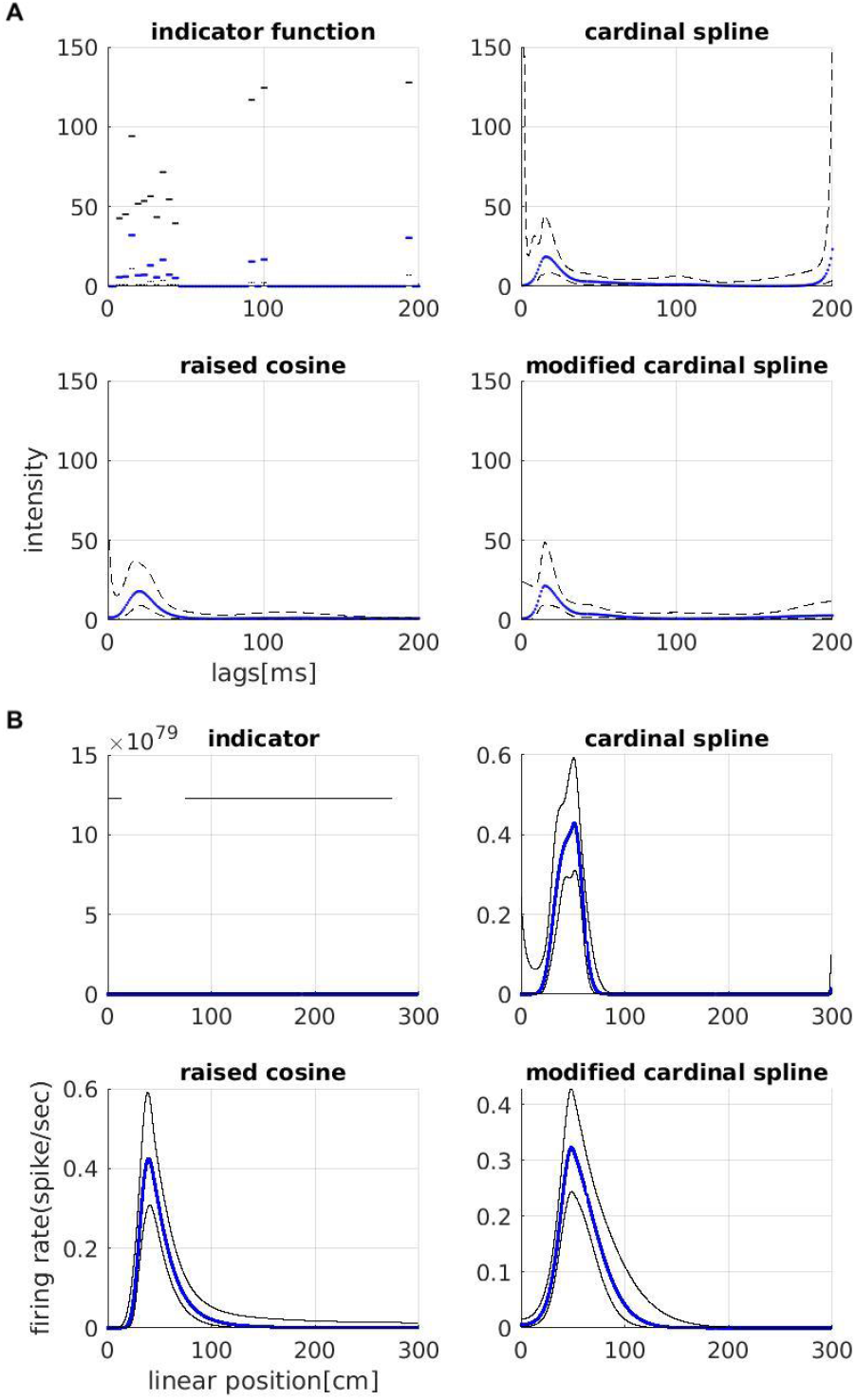
Spatial and temporal components of the neuron’s receptive field. Four classes of basis functions were used in a point process GLM to capture the place field and history dependence of a place cell. A: Firing intensity modulation from each previous spike, going back 200 ms in the past. Intensity modulation is in blue and confidence bounds are in black. B: Spatial component of the firing rate against the linearized position of the animal for the neuron. Blue line represents the place field and confidence bounds are shown in black.

Fig 4B shows the neuron’s place field, estimated using each set of basis functions. The upper left panel shows that the indicator function basis lead to nonsensical confidence intervals (note that the x axis reaches values on the order of 10^79^). This occurs whenever no spike is present over any of the intervals of support for the indicator functions. In that case, the maximum likelihood estimate for the corresponding parameter should go to −∞, at which point, the likelihood will be flat. Numerically, this leads to parameter estimates that are extremely negative and estimated confidence bounds that are enormous. This issue is sometimes called the perfect prediction or complete separation problem. The raised cosine basis does not lead to the perfect prediction problem because each basis element is defined globally rather than locally (ie. it has support over its entire domain). However, globally defined bases can lead to other issues, such as difficulty in interpreting individual model parameters and substantial covariance between those parameters [42, 43]. The raised cosine model fit shows a clear place field at locations between 0 and 100 cm along the track. The cardinal spline basis functions are defined locally (ie. each basis function is defined over a distinct subinterval of its domain), but does not lead to the perfect predictor problem for the model fit for this neuron. The model using unmodified cardinal spline basis functions shows a similar place field structure, but again shows increased uncertainty at the boundaries at positions 0 cm and 300 cm. The model using the modified cardinal spline basis functions shares the advantage of locality with the unmodified basis but corrects the uncertainty at the boundaries.

Fig 5 shows the estimated correlation between the parameter estimates for each basis component used to model history dependence, computed using the inverse Fisher information matrix [10]. For the indicator function basis, there is no off-diagonal correlation, suggesting that each parameter estimate is independent. Each parameter is interpretable on its own - it represents the influence of previous spikes that occur over non-overlapping lags - and its uncertainty is not related to that of other parameters. The correlation matrix for the model with raised cosine basis functions shows a checkerboard pattern, which suggests that the parameter estimates are highly intercorrelated. The influence of a spike at any previous time is determined by a linear combination of all the raised cosine functions, and overestimation of one parameter is likely associated with overestimation or underestimation of all the remaining parameters. This makes it challenging to interpret and perform tests on the influence of past spiking over a particular set of lags (eg. whether a significant refractory period exists). The correlation matrix for the model with unmodified cardinal spline basis functions shows minimal correlation structure for the parameters representing the interior of the domain of lags (parameters 3-5), but substantial correlation between parameters 1 & 2 and between parameters 6 & 7. This makes interpretation and inference about the effects of past spiking more difficult at lags near the boundary. However, the correlation matrix for the model with unmodified cardinal spline basis functions shows minimal correlation between any of the estimated model parameters. Each parameter represents the influence of past spiking over a small set of lags. Parameters representing nearby regions are slightly anticorrelated, and this correlation diminishes for parameters representing more distal regions to each other.

**Fig 5.**
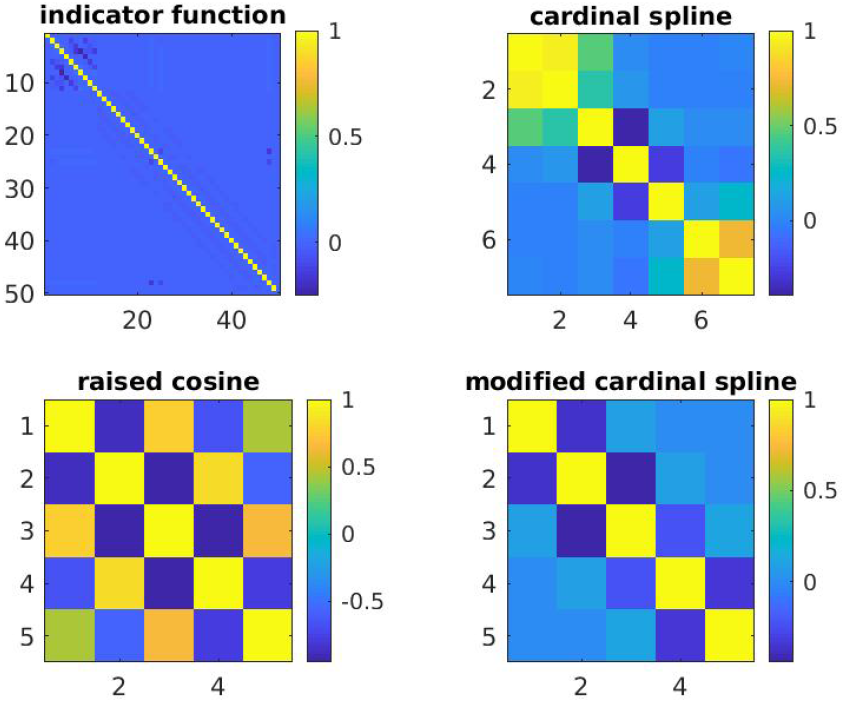
Correlation matrices of the estimated temporal model parameters for each class of basis function. Indicator functions show no off-diagonal correlation between model parameters. Raised cosines show significant correlation structure among all of the parameters, making it difficult to interpret and conduct tests about individual parameters. Unmodified cardinal splines show correlation between parameters near each endpoint, making it difficult to perform tests about parameters that define firing at these endpoints (eg. tests about significance of a refractory period). Modified cardinal splines show minimal correlation between parameters.

While Figures 4 and 5 show the properties of the model fits using different choices of basis functions for a single example neuron, Fig 6 summarizes these properties for a population of 45 neurons in the hippocampus. For the temporal component of the model fits, we plotted the distribution of the confidence interval width ratio (CIWR) between parameters at the boundaries of the set of lags considered over parameters in the interior of this set of lags. The ratio for parameters at the start of the set of lags (lag 0) are shown in blue and the ratio for parameters at the end of the set of lags (lag 200) are shown in red. For models that reflect only the relative uncertainty due to less data around these boundary points, we expect this distribution to be centered between 1 and around 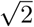. We excluded the indicator basis functions from this analysis because of their tendency to have perfect predictors, which makes the confidence bound ratio less meaningful. For the raised cosine basis functions, the confidence intervals near lag 0 show the expected distribution across all neurons, but the confidence intervals at the largest lags considered are all much smaller than we would expect. This suggests that these basis functions lead to model fits with inappropriately small uncertainty of the influence of spikes at longer lags. Hypothesis tests about these influences are more likely to result in type I errors. For the unmodified cardinal spline basis functions, the distribution for the end boundary is skewed to larger than expected values, suggesting inappropriately large uncertainty at this boundary. Hypothesis tests about these influences are more likely to result in type II errors. For the boundary at t=0, the distribution has modes both at inappropriately small and large confidence interval lengths. We posit that the inappropriately large values arise for the same reason as discussed at the end boundary, while the inappropriately small values likely arise when parameter estimates for the initial spline coefficients become extremely negative leading to confidence bounds whose upper and lower values are both near zero when exponentiated. While the inappropriately large confidence bounds are likely to lead to type II errors, the inappropriately small ones may lead to type I errors. For the modified cardinal spline basis functions, the distributions for both the start and end boundaries fall more in line with our expectations for models reflecting only the relative uncertainty due to the reduction of data near the endpoints. Both distributions are centered between 1 at about 1.5, with few values near 0 or above 3. This suggests that our modifications have been consistently successful at correctly capturing uncertainty near the boundaries.

**Fig 6.**
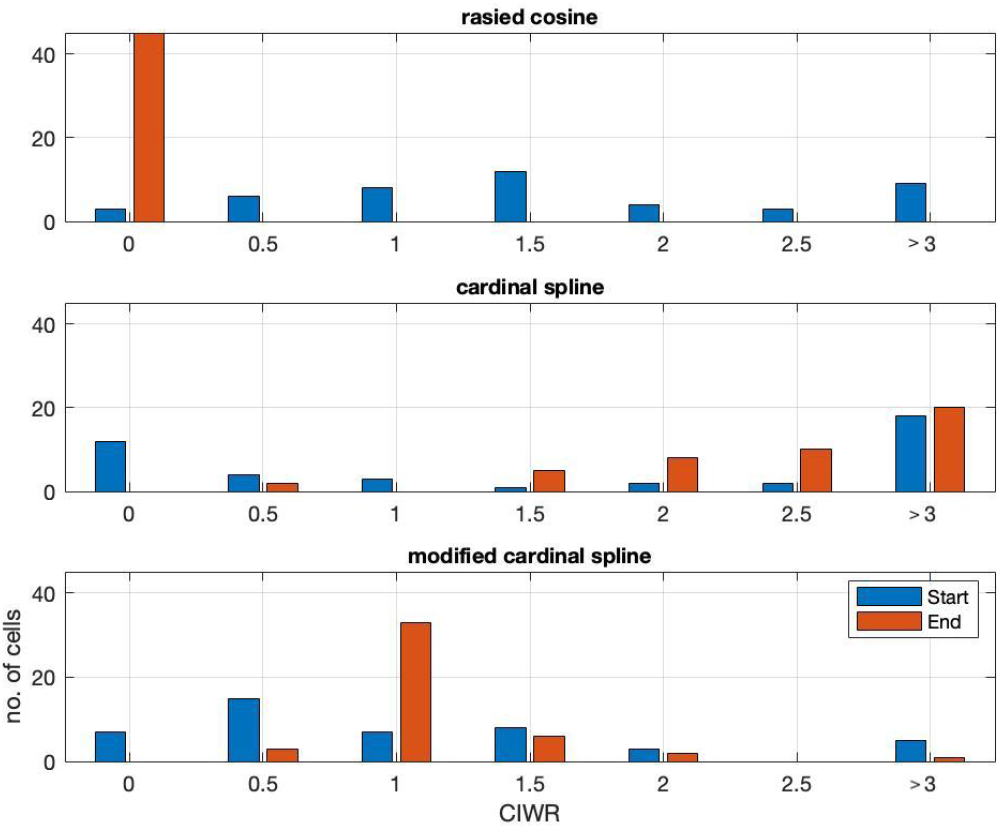
Distribution of relative confidence interval with at endpoints across hippocampal population. Each panel shows for a different set of basis functions the distribution of computed confidence interval width ratio (CIWR) values for each of 45 hippocampal neurons for the temporal component of the model. The distribution for lags near 0 are shown in blue and the distribution for lags near 200 ms are shown in red.

## Discussion

Statistical modeling provides a natural approach for studying neural codes at the level of individual neurons or populations. Point process regression models have been used successfully to capture neural receptive fields from spiking data in a variety of systems and brain areas [1, 9]. Ordinary regression and gamma regression models have also been used to model neural coding properties from imaging and local field data [44, 45]. Since neural receptive fields are rarely fully captured by linear relationships, many successful neural models use basis function expansions to produce flexible, robust, and interpretable model fits. Cardinal splines are a popular choice of basis functions because they are locally defined, smooth, and provide interpretable parameter estimates. However, the need to define additional parameters to control the derivative of the model at its boundaries can lead to a dramatic reduction in statistical power. In this work, we have derived a modification to the cardinal spline basis that increases statistical power at the endpoints by using the assumption that the derivative at the endpoints is zero.

We illustrated the application of this modified basis to the problem of simultaneously estimating the place field and history dependent properties of a set of neurons from the CA1 region of rat hippocampus, and compared the resulting model fits with other commonly used basis functions. These analyses showed that the modified basis maintains the splines’ advantages of smoothness, local definition, and direct interpretability, but has appropriately sized confidence intervals at the endpoints of both the place field and history modulation functions. In addition to improving the statistical power of hypothesis tests about firing around these boundary areas, this makes the confidence intervals arising from neural model fits more visually interpretable. Finally, the fact that parameter estimates representing nearby regions are only slightly correlated and those representing far removed regions are uncorrelated means that meaningful hypothesis tests can be conducted on individual parameters or small subsets of parameters.

The choice of basis provides a trade-off between model flexibility and statistical power. If the goals of an analysis require estimating derivatives of a coding relationship near its endpoints, this modification would not be appropriate. In cases where there are ample data to fit coding relationships near their endpoints, this modification might not be necessary. However, in many cases the assumption of a flat derivative may be a small price for the resulting gain in statistical power. Additionally, goodness-of-fit methods such as AIC analysis [46, 47] and time re-scaling [48, 49] can be used to assess the quality of the flat derivative assumption. However, we note that it may be the case that a model using the unmodified spline basis might lead to a better overall goodness-of-fit than one using the modified basis, where the gain in statistical power is still worth the trade-off.

These methods and applications may be extended in a number of directions. While we developed a particular modification based on the assumption of a flat derivative, other modifications are possible to achieve different levels of trade-off between model flexibility and power. In particular, a different spline modification that uses individual control points at the boundaries to estimate both the conditional intensity and derivative, may be more numerically robust than the unmodified cardinal splines, which use separate control points to estimate the derivative. Additionally, while we focused here on the example of fitting point process models of the receptive fields of spiking neurons, statistical models using standard regression or gamma regression methods have been used to characterize neural coding from imaging and field data as well [44, 45]. Our modified basis functions can be naturally adapted to these model settings. Moreover, here we focused on a simple example that only included two classes of covariates, the animal’s position and the neuron’s own spiking history, in order to highlight issues related to uncertainty in point process GLM estimates near the endpoints of bounded predictors. However, many other influences on these neurons exist (such as the firing of simultaneously recorded interneurons) and it would be interesting to extend this particular analysis by adding additional predictors to more completely capture the spiking structure. However, this is beyond the scope of this work. There are numerous other studies that focus on the broad range of influences that can be fit with point process GLMs [42, 50–54] and that focus on the specific coding properties of these hippocampal neurons [1, 15, 55–57].

For understanding neural coding, statistical models that are flexible, interpretable, and allow for statistically powerful inference are essential. As neuroscience experiments become more complicated, with recordings from larger populations across multiple brain areas and scientific questions involving multivariate relationships during naturalistic tasks, the need for robust modeling tools increases. We believe that the methods developed here, and the associated software tools available at our GitHub repository will provide a valuable addition to the toolbox for neural statistical modeling.

## Acknowledgments

We would like thank Dr. Loren M. Frank at University of California, San Francisco for providing the data and helpful conversations about this work.

